# Accurate single domain scaffolding of three non-overlapping protein epitopes using deep learning

**DOI:** 10.1101/2024.05.07.592871

**Authors:** Karla M. Castro, Joseph L. Watson, Jue Wang, Joshua Southern, Reyhaneh Ayardulabi, Sandrine Georgeon, Stéphane Rosset, David Baker, Bruno E. Correia

## Abstract

*De novo* protein design has seen major success in scaffolding single functional motifs, however, in nature most proteins present multiple functional sites. Here we describe an approach to simultaneously scaffold multiple functional sites in a single domain protein using deep learning. We designed small single domain immunogens, under 130 residues, that simultaneously present three distinct and irregular motifs from respiratory syncytial virus. These motifs together comprise nearly half of the designed proteins, and hence the overall folds are quite unusual with little global similarity to proteins in the PDB. Despite this, X-ray crystal structures confirm the accuracy of presentation of each of the motifs, and the multi-epitope design yields improved cross-reactive titers and neutralizing response compared to a single-epitope immunogen. The successful presentation of three distinct binding surfaces in a small single domain protein highlights the power of generative deep learning methods to solve complex protein design problems.

## Introduction

The “motif scaffolding problem” in *de novo* protein design seeks to recapitulate native protein structural motifs within smaller and simpler protein folds. Motif scaffolding approaches have been used to scaffold metal binding sites, enzyme active sites^1–3^, and to present epitopes inducing broadly neutralizing antibody responses^4–6^ in the hope that presentation of this epitope in the absence of the rest of the antigen may induce more specific and neutralizing antibody responses when administered as a vaccine candidate^6–9^. Despite some success, previous epitope scaffolding approaches have suffered from 1) a lack of potent immune responses elicited by the immunogens, due at least in part to the small relative surface area of epitope to scaffold; and 2) challenges in scaffolding certain epitopes with sufficient accuracy to recapitulate high-affinity antibody binding. Deep-learning based approaches for protein design, which are trained to learn the underlying distribution of possible protein structures^2,3,10^, have been used to scaffold single functional motifs comprising a relatively small fraction of the total residues in the designed proteins.

We reasoned that developing a methodology to accurately scaffold multiple distinct epitopes simultaneously could elicit potent immune responses where a greater fraction of the immune response targets the epitopes versus the scaffold, and provide broader protection than presenting a single epitope alone. Outside of the field of immunogen design, the ability to scaffold multiple functional motifs is relevant to problems such as enzyme design where complex reaction pathways typically depend on multiple functional sites. Specifically, we extended the protein “Inpainting” network RFjoint2 to design de novo scaffolds embedded with up to three distinct epitopes from the respiratory syncytial virus (RSV) fusion protein (RSVF), with arbitrary (non-native) placement of the three epitopes in three-dimensional space.

### *De novo* scaffolds presenting a novel RSVF epitope

We first explored whether deep learning tools are able to solve single epitope-scaffolding problems that have eluded traditional design methods. The site-V epitope of RSVF (RSVFV), consisting of a helix-turn-strand motif, cannot be grafted onto natural scaffolds due to low structural similarity to the Protein Data Bank (PDB) (Fig. <u>S1A</u>). In a previous proof-of-concept, we scaffolded RSVFV into *de novo* proteins using RoseTTAFold motif-constrained hallucination, yielding 3 weak (micromolar) binders to the site-V-specific antibody RSV90 Fab^2^. Here, to attempt to obtain higher-affinity designs, we ran a larger-scale hallucination design campaign and additionally incorporated the ProteinMPNN sequence-design tool to improve stability and solubility^11^. We screened 4,547 designs and 492 bound to RSV90 at 100 nM using a yeast display binding assay. 27 of 39 chosen hits expressed in *E. coli* and we further characterized 4 of the best (RSVFV-1 to RSVFV-4), most structurally diverse designs (Fig. 1A). The designs have good agreement between hallucination models and AF2 predictions, show mixed alpha-beta profiles by circular dichroism (CD) spectroscopy, have minimal unfolding up to 90 °C, and were mostly monomeric by SEC-MALS (Fig. <u>S3</u>). The 4 scaffolds showed K_d_’s between 54 nM-241 nM to RSV90 Fab (Fig. 1B, Supplementary <u>Table 1</u>), within 50-fold of the RSVF trimer (0.9 nM) (Fig. <u>S4</u>) and 20-fold better than the best previous RSVFV design (0.9 μM)^2^. We solved the crystal structure of one of the designs, RSVFV-1, in complex with RSV90 Fab to 2.4 Å resolution. The overall structure of the design was in close agreement with its AF2 prediction (RMSD_backbone_ 1.05 Å) (Fig. 1C), and both the backbone and side chains of the scaffolded RSVFV epitope closely matched their conformations in the RSV90-bound RSVF structure used for design (Motif RMSD: Backbone 0.843 Å, all-atom 1.37 Å) (Fig. 1D-E, Supplementary <u>Table 3</u>). Together, these data show that deep learning methods can scaffold antigenic epitopes with high structural accuracy, and represent a significant improvement over previous RSVFV single epitope scaffolding efforts.

**Figure 1:**
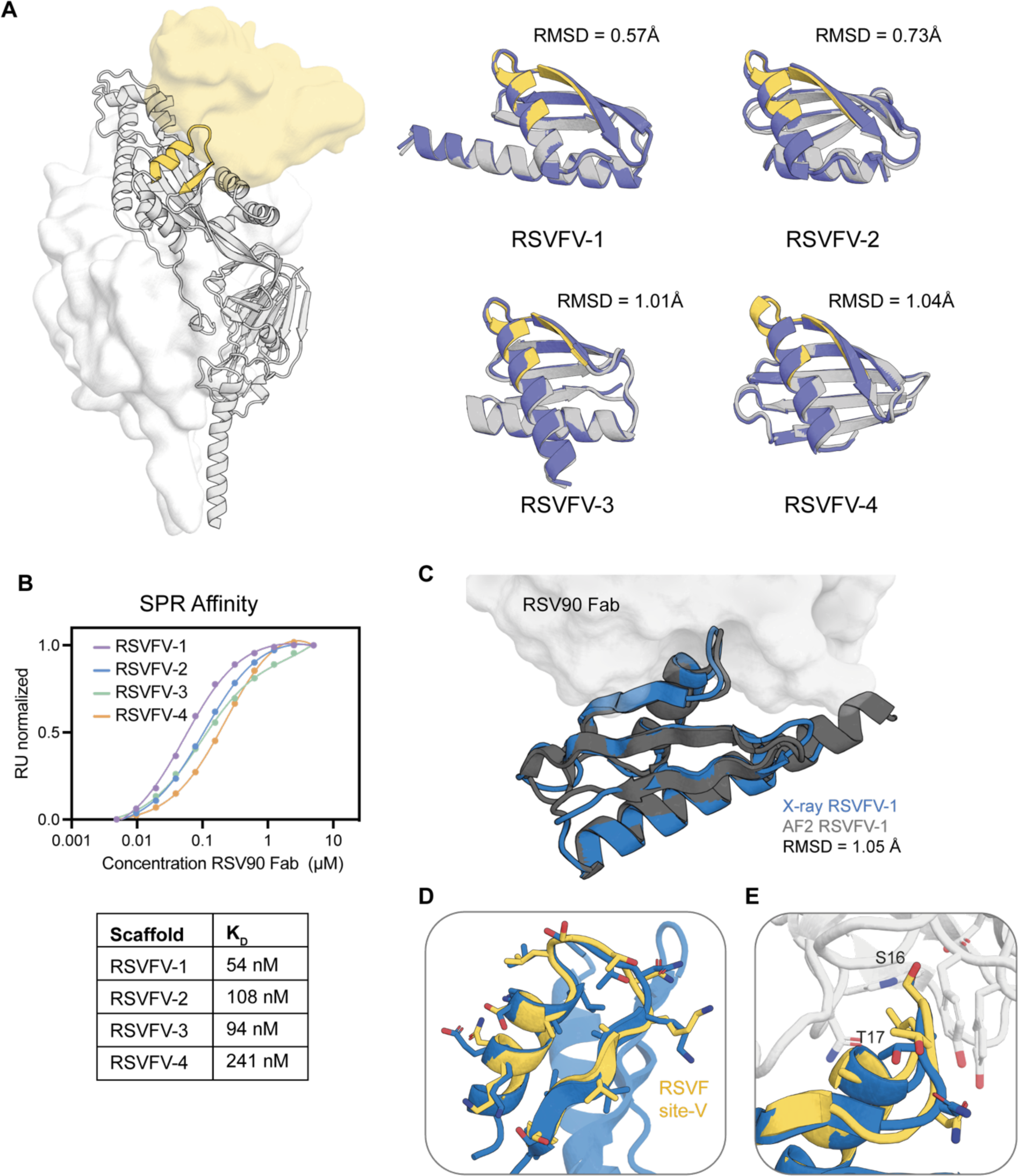
Characterization of RSVFV immunogen candidates. **A**) RSVFV highlighted on the RSVF-trimer bound by a site-V specific antibody (yellow surface). Left panel shows the RoseTTAFold hallucination models (gray) of top candidates scaffolding RSVF site V overlaid on AF2 predictions (blue), all-atom RMSD shown for each model. **B**) SPR steady state affinity measurements for the 4 scaffolds against RSV90 Fab. **C)** Crystal structure of RSVFV-1 in complex with RSV90 Fab closely agrees with the AF2 model (RMSD_backbone_=1.05 Å). **D)** The RSVFV-1 epitope shows high local similarity with the native RSVF site-V structure (RMSD_backbone_ = 0.843 Å). **E)** Superimposition of the major contact residues of RSV90 Fab bound RSVFV-1 superimposed on RSV90 bound RSVF site-V (PDB: 5TPN) (RMSD_all-atom_ = 1.37 Å).

### *De novo* scaffolds incorporating three RSVF neutralizing epitopes

Given the experimental success of deep learning-based protein design methods on scaffolding epitopes considered previously intractable, we next asked whether multiple epitopes of varying structural complexity could be presented simultaneously on a single scaffold, with the aim of generating immunogens with a higher epitope:scaffold surface area and broader neutralization profiles. While computational design approaches have successfully incorporated motifs on *de novo* scaffolds^2,3,12^, few efforts have been aimed towards engineering single domain scaffolds presenting multiple distinct epitopes in non-native orientations^12^.

To design scaffolds embedded with three distinct RSVF epitopes known to elicit neutralizing antibodies (nAbs) (Fig. 2A), we employed the deep learning-based RFjoint2 Inpainting motif scaffolding method, derived from the RF structure prediction network. We provided fragment templates for sites II, IV, and V from RSVF, without specifying their relative geometric positions in the network (Fig. 2B). As these motifs are distinct, their relative positions are not important for eliciting an immune response, and by not pre-specifying their relative geometries a much greater diversity of scaffolds can be generated (Fig. <u>S5</u>). Just as for the individual epitope scaffolding with RF, after the initial RFjoint2 Inpainting design step we used ProteinMPNN^11^ to redesign the sequence of the scaffold and filtered using AF2 as described previously^3^ (Fig. 3A). To eliminate designs in which the antibodies cannot be simultaneously bound to the same scaffold due to steric clashes, we aligned antibodies to the epitopes on the designed scaffold and filtered out designs that were structurally incompatible with the antibody binding mode (Fig. 3B). Because the rigid-body position of the epitopes was not pre-specified, RFjoint2 was able to sample a broad range of different epitope positions (Fig. <u>S5</u>).

**Figure 2:**
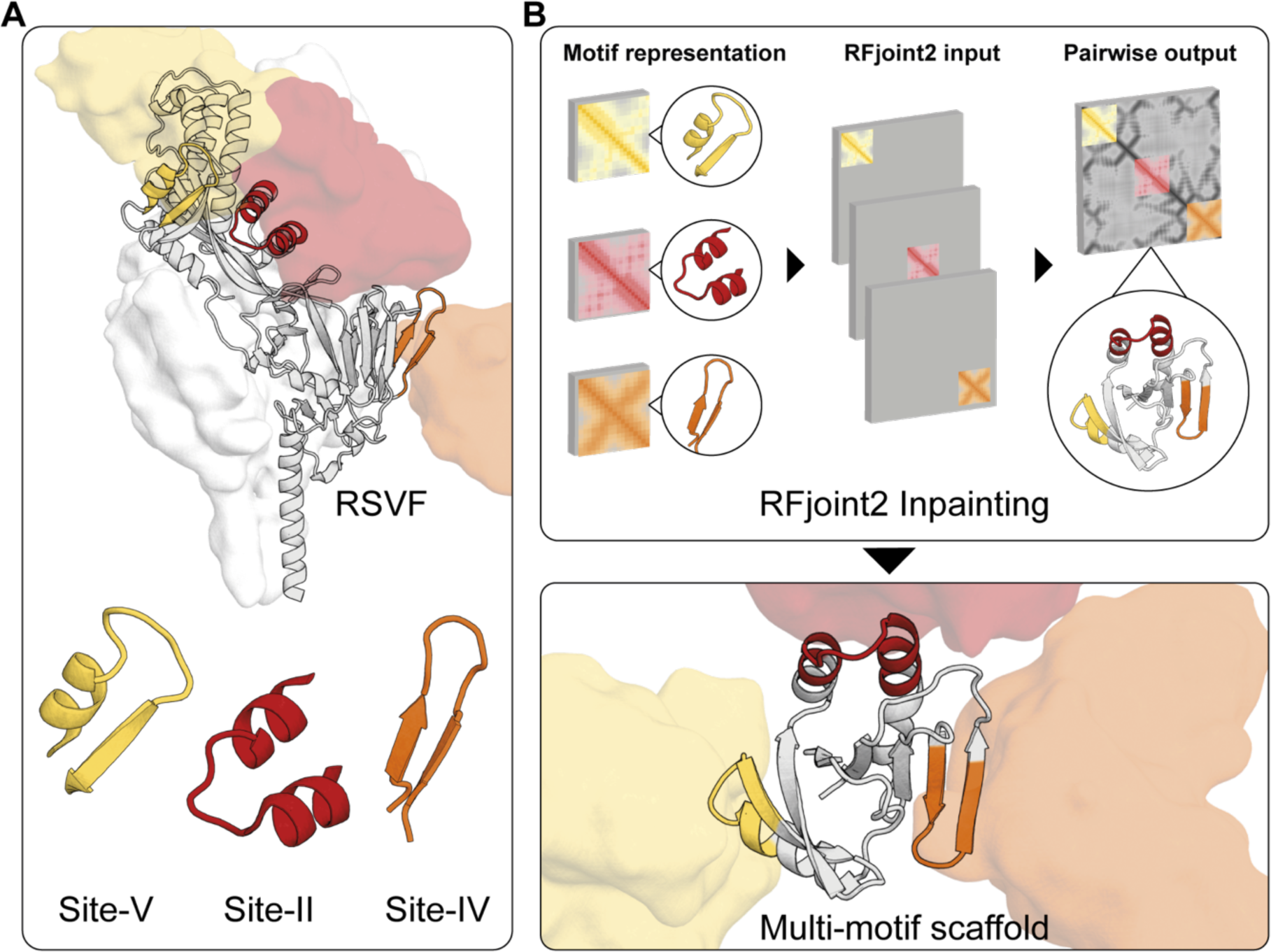
Computational scaffolding of single and multi-epitope immunogens. **A**) Epitopes targeted for single and multi-epitope immunogen design highlighted on RSVF structure (PDB: 5TPN) and the antibody variable domain against site-V (PDB: 5TPN), site-IV (PDB: 3O45), and site-II (PDB: 3IXT) are represented by the corresponding coloured surfaces. **B**) Multi-motif scaffolding utilizes motif templates as input for RFjoint2. Template inputs are invariant to global 3D position, and hence by inputting each motif in a separate template, the relative position of the three motifs is not specified *a priori*, but instead RFjoint2 builds a scaffold (gray protein) that supports the three motifs (colors) in a non-native position with respect to each other.

**Figure 3:**
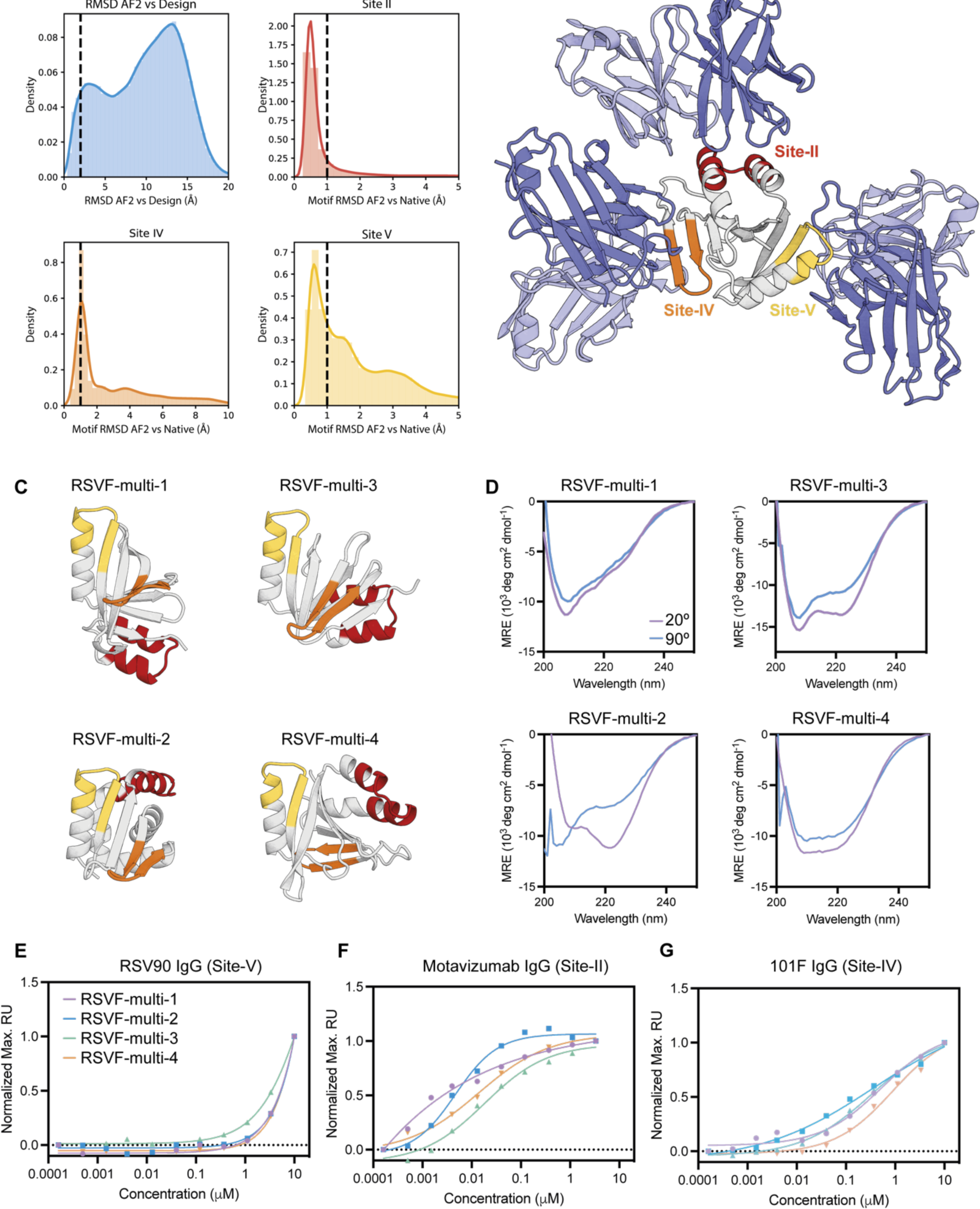
Experimental characterization of multi-epitope immunogen candidates. **A**) *In silico* evaluation. AF2 is used to assess the similarity of predicted structures to the design model and the similarity of each epitope within the prediction to that of the native epitope. Vertical lines: threshold for *in silico* “success”. **B)** Representative design generated with RFjoint2 showing the epitopes remaining accessible to their target antibodies despite this not being explicitly specified during design. The predicted structure of RSVF-multi-4 is aligned to the three known epitope-targeting antibodies (Site II: 3IXT; Site IV: 3O41; Site V: 5TPN). **C**) Predicted structures of top candidate multi-motif scaffolds. **D**) CD spectra at start and end incubation temperatures shown for each scaffold. **E-G**) Normalized SPR steady state affinity measurements for 4 scaffolds for each epitope-specific antibody: RSV90 (site-V), 101F (site-IV), and motavizumab (site-II).

From the filtered multi-epitope designs (RSVF-multi), we selected 32 sequences to screen by yeast surface display. To identify folded scaffolds that successfully recapitulated all three epitopes, we initially subjected the library to limited proteolysis and selection with a site-II specific antibody (motavizumab)^13^. The binding population was then subjected to a second round of selection with a site-IV specific antibody (101F), before a final round of selection with a site-V specific antibody (RSV90) (Fig. <u>S6</u>). From this three-step selection of 32 sequences, we selected 5 high frequency recurring sequences. Four of the five designs expressed well in bacteria. CD spectroscopy of the four multi-epitope designs (RSVF-multi-1, −2, −3, −4) showed mixed alpha-beta profiles in line with the design models, and showed high thermal stability up to 90 °C and mostly monomeric elution profiles (Fig. 3C, S7). All four scaffolds bound to each of the three site-specific antibodies. Quantification of binding affinity for each of the epitopes indicated that both site-IV and site-II specific antibodies bound all four designs with nanomolar affinities (14-47 nM against a site-II-specific antibody, and 343-890 nM against a site-IV-specific antibody), compared to the native RSVF trimer (15 pM to motavizumab Fab and 2 nM to 101F fab) which may be due to the design challenge of combining three epitopes in one scaffold (Fig. 3E-G, <u>S4</u>, <u>Table 1</u>). The low affinity of the designs for the site-V specific antibody likely reflects the structural complexity of this epitope (Fig 3E). However, overall these results demonstrate for the first time that designs displaying multiple multi-functional sites can be generated with relatively high experimental success rates.

### Multi-epitope scaffolds elicit site-specific nAbs in vivo

Following successful validation of our multi-epitope designs *in vitro*, we investigated whether they overcome challenges previously encountered with immunogens where the epitope is only a small fraction of the immunogen’s surface area. We hypothesized that greater immunogenicity might result from immunogens simultaneously displaying multiple different viral epitopes. Previous efforts to recapitulate the RSVF antigenic surface used cocktail vaccine formulations where three neutralizing sites, site-0, site-II, and site-V, were scaffolded and presented on three different protein structures^8^. For these structures, the relevant epitope residues represent no more than 20-39% of the total surface area of each immunogen, likely resulting in a substantial number of antibodies elicited against the host scaffold. By contrast, the multi-epitope designs, by representing the three epitopes on a single scaffold, now represent more than 50% of the total surface area of the immunogen, potentially enabling a superior immune response, specially on the perspective of reducing the decoy surface area presented by the scaffold proteins

For *in vivo* assaying of immune response to the designs, we immunized naive mice with three scaffolds: the single epitope RSVFV-1, and the multi-epitope designs RSVF-multi-1 and RSVF-multi-3. Each scaffold design was incorporated into self-assembling ferritin nanoparticles to improve immunogenicity. Five mice per group were immunized in a homologous boost scheme of three injections of RSVF perfusion trimer or *de novo* nanoparticle immunogens (Fig. 4A). Mice that received an RSVF prime scheme elicited titers and neutralization ability comparable to previous studies (Fig. 4C, 4D)^8,9^. Also, consistent with previous studies of serum neutralization with single epitope scaffolded immunogens, the RSVFV-1 immunogen elicited self-reactive titers, but low levels of pre-fusion RSVF cross-reactive titers, and no detectable neutralizing activity indicating a large proportion of elicited antibodies target the de novo scaffold (Fig. 4B-C)^7–9^. However, for both of the RSVF-multi scaffolds tested we observed moderate levels of pre-fusion RSVF titers, improved from our single epitope immunogen by several orders of magnitude (Fig. 4C). Moreover, RSVF-multi-1 sera showed neutralization activity, albeit still less than pre-fusion RSVF, consistent with previous cocktail vaccine efforts^8^ (Fig. 4D). However, these results confirm that deep learning methods can generate protein-based immunogens with a significantly larger immunogenic surface area compared to single-epitope designs and improved immunogenicity and induction of a physiologically relevant antibody response. These results also validate the previous assumption that the relative surface area of epitope presented is a key determinant of immunogenicity.

**Figure 4:**
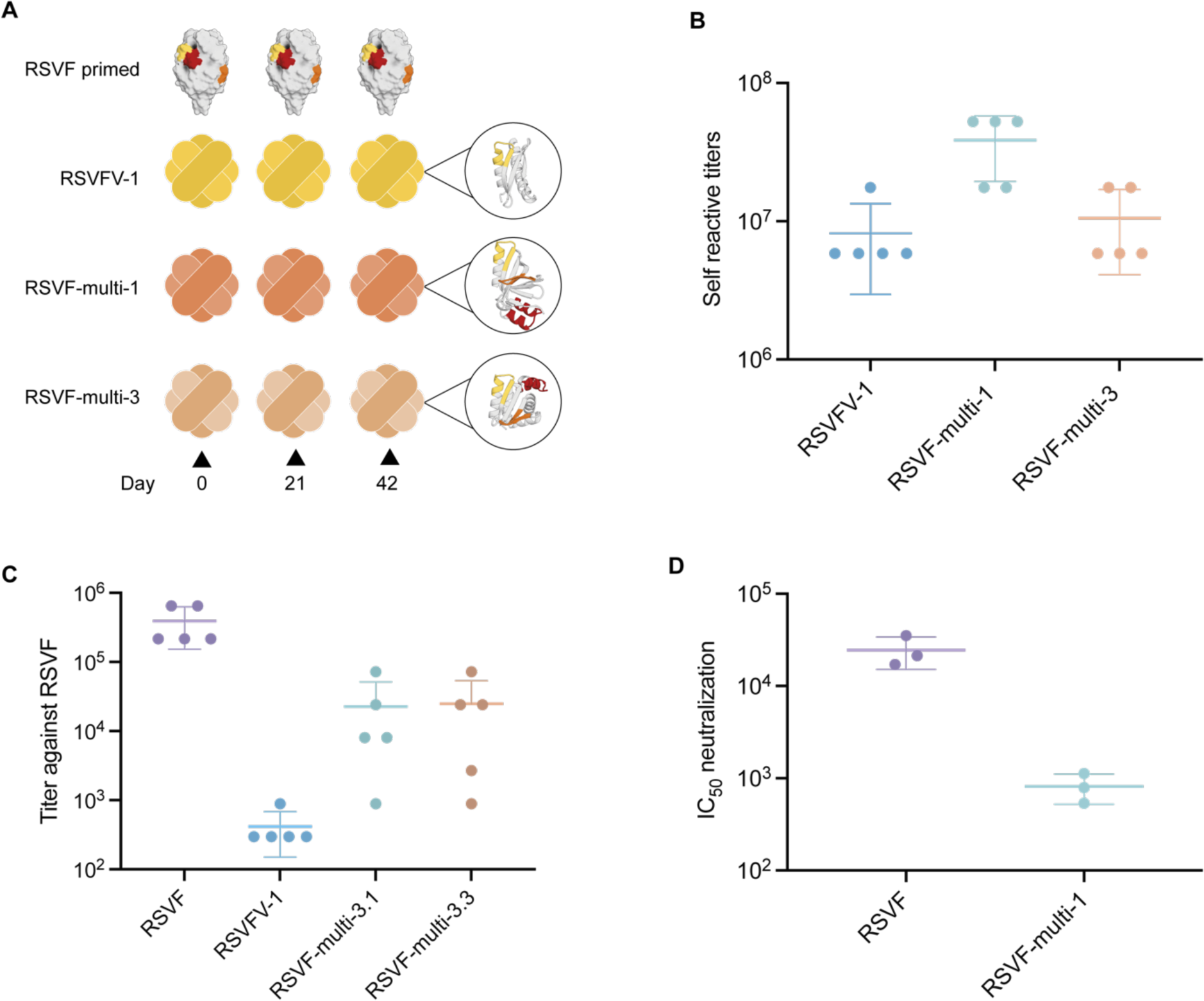
Immunogenicity of single-epitope and multi-epitope immunogens. **A**) Four groups of mice (n=5) were immunized with RSVF followed by adjuvant boosts or a heterologous scheme with the immunogens. **B**) Self-reactive titers of mouse groups immunized with nanoparticle immunogens at day 56 measured by enzyme-linked immunosorbent assay (ELISA). Bars indicate mean and standard deviation. **C**) RSVF reactive titers at day 56. **D)** RSV serum-neutralizing titers from pooled group sera at day 56. Plot indicates calculated neutralization titers from triplicate experiments, with bars indicating mean and standard deviation.

### Multi-motif inpainted designs are structurally accurate

To evaluate the accuracy of the multi-motif designs, we solved the crystal structure of two RSVF-multi immunogens: RSVF-multi-1 in complex with motavizumab Fab and RSVF-multi-4 unbound at 2.91 and 2.3 Å resolution, respectively. Both immunogens closely align to the design models indicating that multi-epitope design can be achieved with a high degree of structural accuracy, RMSD_backbone_ of a approximately 1.4 Å for both designs (Fig. 5A, 5B). We first assessed the structural accuracy of the scaffolded epitopes in RSVF-multi-1 where we observed close mimicry of all three RSVF epitopes (Fig. 5C, Supplementary <u>Table 3</u>). While the structure of RSVF-multi-1 shows slight deviation at the C-terminus of the target site-V backbone structure (RMSD 1.776 Å), the major contact residues S29, T30, N31 (RSVF-multi-1 numbering), and the backbone of K32, are held in the native conformation. The site-IV epitope is composed of a flexible beta-hairpin, and while the majority of the scaffolded epitope accurately mimics the native site-IV epitope, residue 65-66 (RSVF-multi-1 numbering) display deviation from the native structure. Finally, we observe accurate mimicry of the bound site-II epitope in the co-crystallized structure with high agreement of side chain rotamers of conserved scaffolded residues (RMSD_all-atom_ 1.91 Å) to side chain rotamers of the antibody binding interface of native site-II and motavizumab Fab complex (PDB: 3IXT) (Fig. 5A).

**Figure 5:**
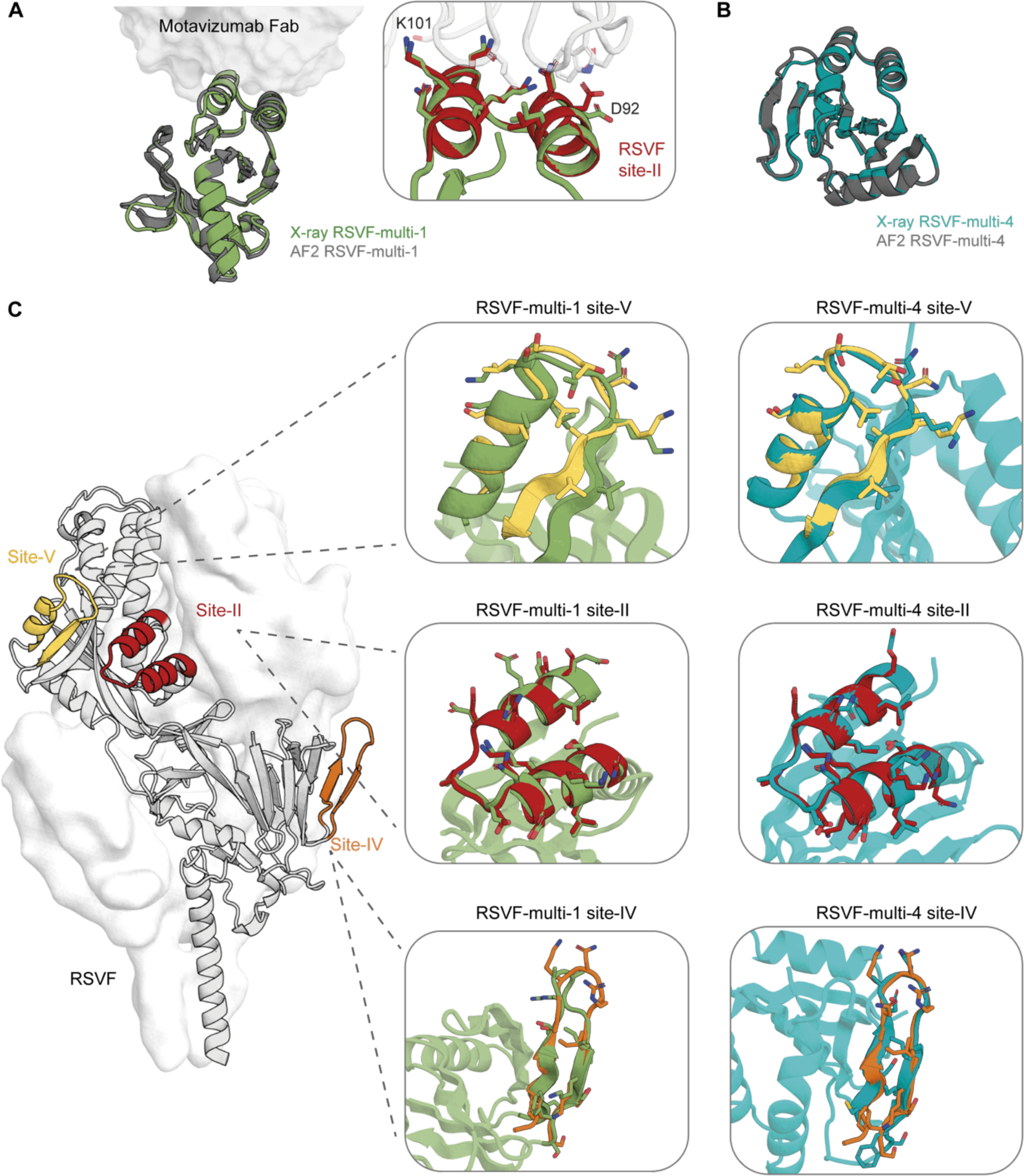
Structural characterization of single-epitope and multi-epitope immunogens. **A**) Crystal structure of RSVF multi-epitope-1 (green) bound to motavizumab superimposed on the AF2 model (gray). Inset panel shows a close up view of major contact residues of RSVF-multi-1 superimposed on motavizumab bound native RSVF site-II (red) (PDB: 3O45) **B**) Crystal structure of RSVF multi-epitope-4 (teal) overlaid on the AF2 model (gray). **C**) Superimposition of crystal structure epitopes to native site-V (yellow), site-II (red) or site-IV (orange) structures of RSVF (PDB: 5TPN).

In the unbound crystal structure of RSVF-multi-4, we observe high similarity of less than 1.2 Å backbone RMSD with all scaffolded epitopes to the native epitope structures (PDB: 5TPN) (Fig. 5C). Similarly to the structure of RSVF-multi-1, the site-IV epitope in one of the well resolved chains shows good agreement with the native epitope in the context of RSVF (PDB: 5TPN) and in the flexible bound epitope peptide (PDB: 3O45). The variation in the site-IV hairpin structure between the four chains indicates the flexibility of the epitope can be stabilized by contacts in the crystal lattice (Fig. <u>S8</u>). Despite this, there is good alignment of the major antibody contact sites, with RMSD of 1.5 - 2 Å (Supplementary <u>Table 2</u>). As the RSVF-multi scaffolds were not co-crystalized with the site-IV specific antibody, the greater deviation observed for this epitope may represent an unbound conformation of this loop region outside its native RSVF context.

Altogether, for all scaffolded epitopes in the structures solved in this paper, we observe close mimicry with RMSD of the backbone and C_β_ to the native epitope below 1 Å in many cases. These results illustrate the capacity of deep learning methods to design accurate *de novo* scaffolds, even whilst scaffolding multiple distinct functional sites

Due to the constraints of scaffolding multiple irregular epitopes the overall folds of the designs are quite unusual. Comparison to the PDB showed that the closest previously known protein structures had TM-scores of 0.46 or lower; 0.5 is generally considered to be the threshold for a new fold. The four RSVF-multi designs are also individually distinct from each other (TM-score < 0.52), indicating that incorporation of all three epitopes did not limit the designable topological space. When clustered by pairwise TM-score, all eight designs are distinct from known CATH topology families (Fig. 6A), with a TM-score to the nearest topology family of 0.39-0.43 (6B). Comparison of the maximum TM-score of our designs to CATH is consistent with the distribution of unique topologies^14^ (Fig. 6C). Taken together, these analyses indicate that the design process created new folds to solve the multi-motif scaffolding problem.

**Figure 6:**
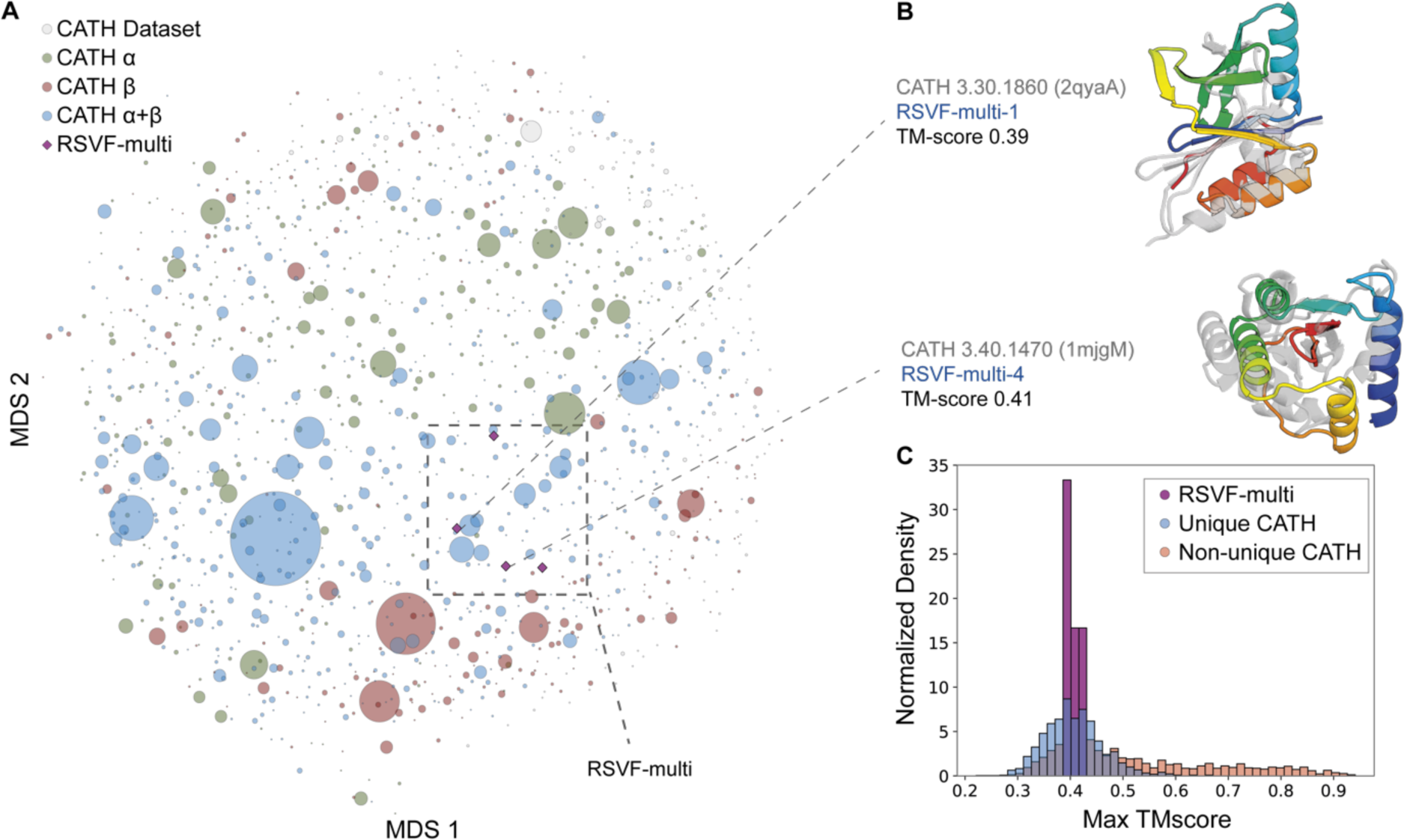
Structural uniqueness of scaffolds presenting single and multi-epitopes. **A)** MDS plot of the pairwise TMscore CATH family representatives, and RSVF-multi designs, coloured by structure class. Circles represent distinct topologies in CATH where the relative size is representative of the number of structures in that family. Purple diamonds represent the four multi-epitope designs. **B)** Structurally confirmed multi-epitope designs (RSVF-multi-1, −4) aligned to a representative of the closest CATH topology family. **C)** Maximum TMscore distribution of generated representations of unique vs. non-unique topologies against CATH topology families. The RSVF-multi designs TMscore to the closest structural homolog fall within the unique topology distribution. RSVF-multi-1 and RSV-multi-3 are shown aligned to a representative of the closest CATH topology family.

## Discussion

The scaffolding of complex motifs has previously been limited by the structural space accessible for design which has restricted *de novo* protein design to a one motif solution per given design problem. In contrast, natural proteins often execute multiple functions with a single chain. Here, we describe the *de novo* design of scaffolds that host up to three structurally distinct motifs in non-native orientations using deep learning approaches. Deep learning-based motif scaffolding offers flexibility in the structures and sequences available to accommodate complex epitopes while requiring significantly less user-input to find compatible scaffolds, greatly improving the accessibility of *de novo* motif-scaffolding design. In contrast to previous single motif design involving large libraries and in some cases in vitro evolution, we have obtained a number of designs binding to all three antibodies using a small number of s designed sequences, a notable step towards the generation of multi-functional designs. Improved affinities could likely be achieved through testing alternative topologies and sequences at higher scale. Moreover, we have shown that RFjoint2 Inpainting can produce a diverse range of topological solutions for multi-motif scaffolding, with different relative motif orientations to a high degree of structural accuracy. We are able to achieve accurate local structural similarity to the native epitope structure and functionality of all three epitopes in novel folds dissimilar to structures in the PDB.

As immunogens, presentation of multiple sites simultaneously could offer improved antigenic surface display. The designed multi-epitope immunogens elicit higher RSVF cross-reactive titers compared to the site-V single epitope immunogen. One of the three-epitope immunogens demonstrates physiologically relevant neutralization titers, implying that the multi-epitope immunogens can mediate an immune response against a larger antigenic surface using a single immunogen compared to using a one-epitope immunogen. Previous work has demonstrated improvement of RSVF titers by using a cocktail vaccine of single-epitope immunogens^8^. Our multi-epitope immunogen can offer an alternative to cocktail vaccines by using a single component, dramatically reducing the cost and improving the speed of production and validation. These multi-epitope designs have a much higher proportion of desirable antigenic surface compared to the single epitope designs, better recapitulating the RSVF antigenic surface on a concise scaffold with reduced potential off-target antibody elicitation.

Beyond immunogens, the ability to incorporate multiple functional sites should be broadly useful for design of new enzymes, sensors, and therapeutic candidates. The structural novelty of the designs coupled with the close agreement with the crystal structures is perhaps the most remarkable illustration to date of the ability of generative deep learning methods to build accurate custom solutions to highly constrained designed problems

## Acknowledgements

We thank A. Hilditch, and M. Pacesa for helpful discussions and comments on the manuscript. We also thank several EPFL facilities: PTPSP (F. Pojer, K. Lau, A.N. Larabi, L. Durrer, and S. Quinche) for protein expression and crystallography support; the phenogenomics center for support with mouse experiments; the flow cytometry core facility; the gene expression core facility for support with next-generation sequencing; and SCITAS for support in high performance computing.

## Author Contribution

K.M.C., J.L.W., J.W., D.B. and B.E.C. conceived the work and designed the experiments. J.L.W. and J.W. performed the computational design and analysis. K.M.C., A.R., S.R. performed the experimental characterizations. K.C. solved the crystal structures. J.S. performed structural and sequence diversity analysis. K.M.C. and S.G. contributed to the design and planning of animal studies, and sera analysis. K.M.C., J.L.W., J.W., D.B., and B.E.C. wrote and edited the manuscript with input from all authors.

## Data Availability

Structures have been deposited in the Protein Data Bank under accession codes 9F91 (RSVFV-1 in complex with RSV90 Fab), 9F90 (RSVF-multi-1 in complex with motavizumab Fab), and 9F8Y (RSVF-multi-4). Plasmid sequences are available in Supplementary <u>Table 4</u>. The plasmids of the designed proteins are available from the authors under a material transfer agreement with the Ecole Polytechnique Fédérale de Lausanne (EPFL). All code used for this study will be made available.

## Materials and Methods

### Design of site-V presenting topologies

We designed RSV-F site V (PDB 5TPN, residues 163-181) scaffolds using a published RoseTTAFold hallucination method^2^. First, 10,000 designs were hallucinated by 600 steps of gradient descent with a repulsive loss (*α*=3.5 Å, weight = 2) and a radius of gyration loss (threshold = 16 Å, weight = 1). Half of the designs used a larger definition of site V motif (chain A residues 163-191) to include an additional beta-strand to potentially improve recapitulation of the main motif. Motif residue amino acids were allowed to change except for those in contact with the antibody (165-166, 169-178, 180, 188), which were fixed to wildtype. 577 designs with AF pLDDT > 75, AF motif RMSD < 1.2 Å and radius of gyration < 16 Å were used as “seeds” for additional iterations of hallucination. Starting from each seed, multiple trajectories of 300-1000 Markov Chain Monte Carlo (MCMC) steps were run with surface nonpolar (weight 1) and net charge (target charge −7, weight 0.02) losses. Designs were filtered on AF pLDDT, AF motif RMSD, and SAP score and used to seed additional rounds of MCMC. Finally, designs were subjected to 20-100 steps of MCMC in the presence of the target antibody to eliminate side-chain clashes with the antibody from residues outside the epitope motif.

A total of 47,498 designs were generated over 14 rounds of hallucination; these were filtered down to 5,717 designs by the following metrics: AF pLDDT > 82, AF motif RMSD < 1.0 Å, hallucination motif RMSD < 1.0 Å, AF-hallucination whole-protein RMSD < 2.0 Å, radius of gyration < 16 Å, SAP score < 35, net charge < −5, Rosetta score/residue < −2.5, Rosetta ddG < - 10. “Motif RMSD” refers to the RMSD over backbone atoms (N, C-alpha, C) between either the AF prediction or the hallucination (RoseTTAFold-predicted structure on the last step of gradient descent or MCMC) and the input crystal structure motif. Rosetta ddG was calculated after superimposing the design on the native motif in complex with hRSV90 antibody and minimizing side chains in Rosetta. This set of designs was further reduced to 3,072 by clustering at 90% sequence identity using MMseqs2^15^ and keeping one design from each cluster with the lowest mean of AF and hallucination motif RMSDs. This comprised the “hallucination-only” subset of the experimentally tested designs.

The ProteinMPNN subset of designs were created by clustering the final hallucinations at 70% sequence identity, to 911 designs, combining with 230 high-scoring seed hallucinations (from gradient descent hallucination before iteration), and inputting their AlphaFold2 models to ProteinMPNN for sequence redesign with temperature 0.1 and fixing interface residue amino acids. The target antibody was not included in ProteinMPNN design. 8 sequences were generated from each input backbone, yielding 9128 designs. 592 designs were chosen for testing using the same metric thresholds as above but slightly stricter on motifs: AF motif RMSD < 0.8 Å, hallucination motif RMSD < 0.8 Å. Finally, high-scoring ProteinMPNN designs were subjected to Rosetta-based mutation of surface residues to adjust net charge to −7, as this has been reported to improve the success rate of designed protein binders^16^, filtered using the same thresholds as above, and clustered at 90% sequence identity to obtain 883 designs for testing. The yeast display screen shows that ProteinMPNN without adjusting charge was the most successful design method, with a success rate at 10-fold binding enrichment of 33.7%, compared to 3.3% for hallucination-only and 21.7% for ProteinMPNN and charge adjustment (Fig. <u>S2A</u>).

When selecting designs for experimental testing, we filtered on *in silico* metrics using thresholds that were chosen by intuition. After experimental testing, we examined the receiver operating characteristic area under the curve (AUC) between each metric and binding success at 1-fold or 10-fold enrichment between binding and non-binding yeast populations (Fig. <u>S2B</u>-C)^16,17^. AUC was relatively high (between 0.7 and 0.8) for several metrics when considering all designs, but within each of the 3 design methods, AUC was much lower and usually close to random (0.5). This suggests that most of the differences in success rate were driven by the design method, and that more-stringent filtering on the metrics we used would not have increased success further.

### Designing multi-epitope scaffolds without specifying inter-epitope rigid body orientation

We sought to develop a design methodology to permit the scaffolding of multiple epitopes into a single designed protein. Importantly, because each individual epitope can be considered its own unit, we want a methodology whereby the rigid-body orientation between the epitopes can be sampled/chosen during design. In other words, we do not want to keep the native rigid-body orientation, nor do we want to have to pre-specify this rigid body orientation.

We therefore developed an approach using RFjoint2 Inpainting, taking advantage of its ability to accept multiple independent “templates” (inherited from the original RF model from which it is fine-tuned). During protein structure prediction, RF uses any available homologous structures to aid it in structure prediction. Importantly, there can be multiple templates, which align to distinct and non-overlapping regions of the query sequence. Hence, these multiple templates are provided to RF in a rigid-body orientation-independent manner, such that the rigid body orientation between the two templated regions is not specified/fixed during prediction (because it is unknown). This invariance comes from the pairwise representation with which templates are provided to RF. Within each template, the pairwise residue-residue distances and dihedral angles^18^ are provided to the network. The pairwise distances/angles between templates is therefore not provided to RF.

We exploited this in RFjoint2 Inpainting, to develop a method whereby individual epitopes could be provided as individual templates to the network. In other words, the three RSVF epitopes we scaffolded (sites II, IV and V) were each provided to the network as their own distinct template input. Thus RFjoint2 Inpainting was able to see each epitope’s internal structure, but not the rigid body orientation between them. RFjoint2 therefore had to simultaneously build a scaffold (with the specified masked residues that connected the epitopes) and “choose” the rigid body orientation between the three epitopes. The ability to perform this task is inherent to RFjoint2 Inpainting, and did not require further training.

### Designing multi-epitope scaffolds scaffolding RSV sites II, IV and V

#### Backbone generation with RFjoint2

The structure of sites II, IV and V were extracted from PDB: 4JHW. The three sites were defined as:

| Site | Residues |
| --- | --- |
| II | F255-277 |
| IV | F420-436 |
| V | F163-181 |

Because the rigid-body orientation between epitopes is not specified, there is no guarantee that designs will leave each epitope accessible to the target antibody. Pilot experiments demonstrated that multi-epitope designs often had detrimental clashes with the 101F Site IV antibody (from PDB: 3O41). To prevent this, we therefore additionally included residues H1-116 of the 101F Fab (from PDB: 3O41) in the Site IV input template.

To generate a diverse set of designs, the order of the three epitopes in the designs was enumerated, and the lengths connecting each epitope was systematically sampled (10-20 amino acids at the N- and C-terminus, and connecting the epitopes). A total of 17475 designs were made (approximately 20% of the total possible enumerated design inputs with these parameters). In all cases, the “template confidence” (i.e. how confident RFjoint2 should be in the internal structure of each input epitope) was set to 1.

#### Sequence design with ProteinMPNN

Although RFjoint2 Inpainting simultaneously designs a structure and sequence, the ability of ProteinMPNN to very rapidly design multiple unique sequences for a given protein backbone renders it preferable over the single sequence that RFjoint2 produces for a given protein backbone. We therefore used ProteinMPNN for the sequence design step of the design pipeline.

To reduce the compute burden, we pre-filtered RFjoint2 outputs prior to downstream sequence design and filtering. Specifically, we filtered both on the internal confidence metric (pLDDT) within RFjoint2 (> 0.6), as well as on the “compactness” of the outputs (radius of gyration, aspect ratio and the maximum distance between any two residues).

With these filtered backbones, we designed 16 sequences for each backbone with ProteinMPNN^11^, with default parameters. Because not all of the residues within each epitope are actually involved in binding the target antibodies, we allowed ProteinMPNN to redesign the following residues:

| Site | Residues permitted for redesign |
| --- | --- |
| II | F255-261,F264-267,F270,F273-274,F277 |
| IV | None |
| V | F163-168,F171-172, F177, F179-181 |

#### Re-predicting the structure with AlphaFold2

Following numerous other works^2,3,19^, we predict the structure of the sequences out of ProteinMPNN using AlphaFold2 (AF2), to determine the similarity of the AF2 prediction and the designed protein backbone (the “self-consistency”). We therefore predicted the structure of the sequences using AF2_model_4_ptm, with 3 recycles.

#### Calculating interface metrics with Rosetta

In addition to filtering on AF2 self-consistency, we filter on the quality of the interface with each target antibody, as assessed by Rosetta^20^. For this, we aligned the AF2 predictions of each design onto the original antibody-epitope structure (Site II, PDB: 3IXT; Site IV, PDB: 3O41; Site V, PDB: 5TPN). Following Rosetta FastRelax^20^, we calculated the predicted ddg of the interface.

#### Filtering designs for experimental characterisation

For testing, we filtered designs using the following metrics, which largely follow previous work^2,3,16^.

- Epitope RMSD, AF2 vs Native < 1Å
- Overall RMSD, AF2 vs design model < 2Å
- AF2 pLDDT > 80
- Rosetta ddg for each epitope aligned to its respective antibody < −10
- Rosetta SAP score < 40

#### Yeast library preparation

Pools of DNA oligonucleotides computationally designed sequences were purchased through Twist Biosciences. The DNA 3′ and 5′ ends were constant, for the amplification and attachment of overhangs for yeast transformation by PCR. Oligo pools were resuspended in H2O to 50 ng/uL and amplified by PCR (55 °C annealing for 30 sec, 72 °C extension time for 1 min, 25 cycles) The PCR also allowed for attachment of homologous overhangs for yeast transformation. The PCR product was desalted and transformed together with linearized pCTcon2 vector or pNTA_V5 vector as previously described^21^ into EBY-100 yeast strain with a transformation efficiency of at least 10^6^.

#### Yeast surface display

The transformed cells were passaged twice in SDCAA medium with pen/strep before induction in SGAA medium overnight at 30 °C. Induced cells were pelleted by centrifugation (3000 x g, 3 minutes). Pellets corresponding to 2 mL at OD600 of 1 were washed once and resuspended 250 uL in TBS (20 mM Tris pH 8.0, 150 mM NaCal). Chymotrypsin was diluted in 250uL at 2x final concentration. Final concentrations of chymotrypsin were 0.01 uM (pCTcon2 vector) or 0.1 uM trypsin (pNTA-V5 vector). Chymotrypsin was added to the resuspended yeast and incubated at room temperature for 5 minutes. The reaction was quenched in cold PBS + 2 % BSA. Cells were pelleted and washed in cold wash buffer (PBS + 0.05% BSA) three times. Cells were labelled with 1 µM of the target (RSV90 Fab for RSVFV library, motavizumab IgG for the RSVF-multi library) at 4 °C for 2 hours. Cells were washed twice with wash buffer and then incubated with FITC-conjugated anti-HA and PE-conjugated anti-human Fc (BioLegend, #342303) or PE-conjugated anti-Fab (Thermo Scientific, #MA1-10377) for an additional 30 min. Cells were washed and sorted using a SONY SH800 flow cytometer in “ultra-purity” mode. Cells were grown in SDCAA media. The RSVF-V library was subject to a second sort at 0.1 µM RSV90 Fab. Following the sorts, cells were plated on SDCAA agar and more than 40 single colonies were sequenced to obtain 19 candidates for biochemical characterization. The RSV multi-epitope library was subject to a second sort at 1 µM101F IgG and third sort at 1 µM RSV90 IgG to screen against all three antibodies. Cells from the third sort were plated and 40 single colonies were sequenced for which the converged 5 sequences were selected as candidates for biochemical characterization.

#### MiSeq sequencing

After sorting, cells were passaged once in SDCAA medium with pen/strep. The cells were pelleted and lysed for DNA extraction using Zymoprep Yeast Plasmid Miniprep II (Zymo Research) using the manufacturers instructions. The extracted DNA was amplified using plasmid-specific primers with standard Illumina sequencing adapters were attached with overhang PCR in reaction I. Illumina sequencing primers with unique barcodes were added in PCR reaction II. PCR products from both reactions were cleaned using AMPure XP Bead-Based Reagent (Beckman Coulter). Eluted DNA was validated by Fragment analyzer prior to submission for Illumina MiSeq by the Gene Expression Core Facility (EFPL, Switzerland). 500 paired end cycles were run with approximately 2 million reads/sample. Sequences were trimmed and translated in the correct reading frame for bioinformatic analysis, and fold enrichment was computed for each sequence as the fraction counts in the binding population over the non-binding population. Binding designs were considered if a 10-fold enrichment in binding versus non-binding populations was observed. Designs for biochemical characterization were selected among reoccuring sequences from MiSeq in combination with single colony sequencing. Designs whose models were structurally diverse were selected.

#### Surface plasmon resonance

SPR measurements were performed on a Biacore 8K (GE Healthcare) in 10 mM HEPES pH 7.4, 150 mM NaCl, 3 mM EDTA, 0.005% v/v Surfactant P20 (GE Healthcare). Scaffolds and RSVF trimer or IgG proteins were immobilized on a CM5 chip (GE Healthcare # 29104988) by amine coupling. Approximately 1000 response units (RU) of protein were immobilized, and antibody Fab or designed monomeric proteins were injected as analyte in 3 or 6-fold serial dilutions. The flow rate was 30 μl/min for a contact time of 120 s followed by 600 s dissociation time. After each injection, the surface was regenerated using 0.1 M glycine at pH 3.0. Data were fitted using 1:1 Langmuir binding model within the Biacore 8K analysis software (GE Healthcare #29310604).

#### Circular dichroism

Circular dichroism spectra were measured using a Chirascan instrument in a 1-mm path-length cuvette. The protein samples were prepared in 10 mM sodium phosphate buffer at a protein concentration of ∼30 μM. Wavelengths between 195 nm and 250 nm were recorded with a scanning speed of 20 nm min-1 and a response time of 0.125 secs All spectra were averaged two times and corrected for buffer absorpton. Temperature ramping melts were performed from 20 to 90 °C with an increment of 2 °C/min or 5 °C/min. Thermal denaturation curves were determined by the change of elliptcity mnmum at 220 nm.

#### Immunogen expression and purification

Genes encoding scaffolds were purchased as DNA fragments from Twist Bioscience and were cloned into pET11b bacterial expression vector. Plasmids were transformed into E. coli BL21 (DE3). Following overnight preculture growth, cultures were grown in TB autoinduction media at 37 C to OD600 0.7 then transferred to 18 C for 16 hours. Cells were harvested and pellets were resuspended in lysis buffer (50 mM Tris, pH 7.5, 500 mM NaCl, 5% Glycerol, 1 mg/ml lysozyme, 1 mM PMSF, and 1 μg/ml DNase) supplemented with 1x CellLytic B Cell Lysis Reagent (Merk). Chemical lysis was performed rotating at 4 C for 1 hour. Lysates were clarified by centrifugation (48,000g, 20 minutes) and purified by Ni-NTA affinity chromatography eluting with 10 mM Tris, 500 mM NaCl, and 300 mM Imidazole (pH 7.5) and subsequent size exclusion chromatography on a HiLoad 16/600 Superdex 75 column (GE Healthcare) in PBS buffer.

#### RSVF protein expression and purification

The pre-fusion RSVF sc9-10 DS-Cav1 A149C Y458C S46G E92D S215P K465Q thermostabilized variant was previously cloned and annotated as DS2 ^9,22^. The plasmid was transfected in HEK293-F cells and cultured in FreeStyle medium. Supernatants were harvested after 7 days and purified by Ni-NTA affinity chromatography eluting using 10 mM Tris, 500 mM NaCl, and 300 mM Imidazole (pH 7.5). The eluted protein was further purified on a StrepTrap HP affinity column (GE Healthcare) eluted using 10 mM Tris, 150 mM NaCl and 20 mM desthiobiotin (pH 8) (Sigma). Final purification by size exclusion chromatography in PBS (pH 7.4) on a Superdex 200 Increase 10/300 GL column (GE Healthcare).

#### Nanoparticle-linked immunogens expression and purification

The target scaffold genes for RSVFV-1, RSVF-multi-1 and RSVF-multi-3 were fused upstream of a gene encoding Helicobacter pylori ferritin (GenBank ID: QAB33511.1) and N-terminal 6x His Tag. The fusion consisted of a GGSGGSGG, GNGSGGNGSGAEAAAKEAAAKAGNGSGGNGS, or GTGGSGGSGG linker, respectively. The fused genes were cloned into the pHLsec vector for mammalian expression. Ferritin-linked immunogens were transfected in HEK-293T cells. The supernatant was harvested after 6 days and purified using Ni-NTA affinity chromatography and size exclusion on a Superose 6 increase 10/300 GL column (GE).

#### Antibody IgG and Fab protein expression and purification

Heavy and light chain DNA sequences of antibody fragments (Fab) were purchased from Twist Bioscience and cloned individually into the pHLsec mammalian expression vector (Addgene, #99845) using Gibson assembly. The heavy chain sequence was additionally cloned into pHLsec-Fc containing the human C_H_2 and C_H_3 region. HEK-293T cells were transfected with a 1:3 ratio of heavy to light chain. Supernatants were collected after 6 days and purified using a 5-ml HiTrap Protein A HP column (GE Healthcare) for IgG expression and 5-ml kappa-select column (GE Healthcare) for Fab purification. IgG or Fabs were eluted with 0.1 M glycine buffer (pH 2.7), immediately neutralized by 1 M Tris buffer (pH 9), and further purified by size exclusion chromatography on a Superdex 200 Increase 10/300 GL column (GE Healthcare) in PBS.

#### Mouse immunizations

All animal experiments were approved by the Vaud Veterinary Cantonal Authorities in accordance with Swiss regulations of animal welfare (VD3808). Female Balb/c mice at 5-weeks old were obtained from Janvier labs and acclimatized for one week. Immunogens were mixed with equal volumes of adjuvant (Alhydrogel, Invivogen) and incubated for 1 hour on ice. Mice were injected subcutaneously with 100 uL of vaccine formulated with 5 ug of immunogen. Immunizations were performed on days 0, 21, and 42. Blood samples were collected on days 0, 14, and 35. Mice were euthanized on day 56 and blood collection was performed by cardiac puncture.

#### ELISA

Purified recombinant RSVF trimer was coated on Ninety-six well plates (Nunc MediSorp, Thermo Scientific) overnight at 4 C with 0.5 ug/mL in coating buffer (phosphate buffered saline (PBS) pH 7.4). Plates were washed three times at each step with wash buffer (PBS + 0.05% Tween-20 (PBS-T)) Wells were blocked with blocking buffer (PBS-T with 5% skim milk (Sigma)) for 1h at room temperature. Three-fold serial dilutions were prepared in assay buffer (PBS-T + 1 % Bovine Serum Albumin) and were added to the plates and incubated at room temperature for 2 hours. Secondary incubation with anti-mouse (abcam, #99617) HRP-conjugated secondary antibody diluted 1:1,500 was added and incubated for 1 hour at room temperature. Plates were developed by adding 100 uL TMB substrate (Thermo Scientific). The reaction was stopped using and equal volume of 0.5 M HCl. The absorbance at 450nm was measured on a Tecan Safire 2 plate reader. The titer was determined as the reciprocal of the serum dilution which resulted in a signal two-fold above background.

#### Neutralization assay

RSV neutralization was performed as previously described^8^. Hep2G cells were seeded in a Corning 96-well tissue culture plates (Sigma) at 40,000 cells/well overnight at 37 C at 5% CO2. Serial dilutions of heat-inactivated sera were prepared in M0 media (EMEM without phenol red + L-glutamine 2 mM + Pénicilline 100 IU/mL + Streptomycin 100 µg/mL) and incubated with 100 pfu/well RSV-Luc (RSV Long strain carrying a luciferase gene) for 1 hour at 37 C. The sera with virus was added to the cell monolayer and incubated for 48 hours at 37 C. Cells were lysed in 100 uL lysis buffer (31.25 mM Tris pH 7.9, 10 mM MgCl_2_, 1.25% Triton X-100, 18.75% glycerol) for 10 minutes at room temperature. 50 uL lysate was transferred to a white ninety-six well plate (Grenier). An equal volume of assay buffer (lysis buffer + 1 mM DTT, 1.1 ug/mL luciferin (Sigma, #L-6882), and 2 mM ATP (Sigma, #A3377)) were added and luminescence was immediately measured on a Tecan Spark plate reader. The neutralization curve was plotted and fitted using the GraphPad variable slope fitting model, weighted by 1/Y^2^.

#### X-ray crystallization and structural determination co-crystallization of complex RSV90 Fab with RSVFV-1

##### Co-crystallization

The RSVF-V-1 and RSV90 Fab were purified by size exclusion chromatography using a Superdex200 26 600 (GE Healthcare) equilibrated in 10 mM HEPES pH 7.5 and 150 mM NaCl and concentrated to ∼20 mg/ml (Amicon Ultra-15, MWCO 3,000). The immunogen and fab were mixed in equimolar quantities and incubated at 4 C for 1 hour. Crystals were grown at 18 C using the sitting-drop vapor-diffusion method in drops containing 1 μl of protein complex and 1 μl of reservoir solution containing 0.1 M HEPES pH 7.5, and 30 % (v/v) PEG Smear Low. Crystals appearing after 5 days were crushed and used for seeding. Crystals were grown at 18 C in drops containing 1 μl of protein complex, 1 μl of reservoir solution, and 1 uL seeding crystal solution. Crystals appeared in condition solution containing 0.1 M MES pH 6.5 and 20 % (v/v) PEG Smear High. For cryoprotection, crystals were briefly immersed in mother liquor containing 20% ethylene glycol.

##### Data collection and structural determination

Diffraction data were recorded with X06SA (PXI) at the Swiss Light Source, Switzerland. The diffraction data were integrated and processed to 2.43 Å by AutoProc with a high-resolution cut at I/σ=1 applied. The crystals belonged to space group P 21 21 2. The structure was determined by the molecular replacement using the Phaser module in the Phenix program^23^. The searching of the initial phase was performed by using the RSV90 structure (PDB 5TPN) and the AF2 model of RSVFV-1 as a search model. One fab copy contained loops opposite the interaction interface which could not be built. Manual model building was performed using Coot ^24^, and automated refinement using Phenix Refine. The final refinement statistics are summarized in Supplementary <u>Table 2</u>.

#### X-ray crystallization and structural determination co-crystallization of complex motavizumab Fab with RSVF-multi-1

##### Co-crystallization

The RSVF RSVF-multi-1 immunogen and motavizumab Fab were purified by size exclusion chromatography using a Superdex200 26 600 (GE Healthcare) equilibrated in 10 mM HEPES pH 7.5 and 150 mM NaCl and concentrated to ∼20 mg/ml (Amicon Ultra-15, MWCO 3,000). The immunogen and fab were mixed in equimolar quantities and incubated at 4 C for 1 hour. Crystals were grown at 18 C using the sitting-drop vapor-diffusion method in drops containing 1 μl of protein complex and 1 μl of reservoir solution containing 0.2 M potassium thiocyanate, 0.1 M sodium cacodylate pH 6.5, and 10 % (v/v) PEG 1000.

##### Data collection and structural determination

Diffraction data were recorded with ID30B at the European Synchrotron Radiation Facility, Switzerland. The diffraction data were integrated and processed to 2.692 Å by AutoProc with a high-resolution cut at I/σ=1 applied. The crystals belonged to space group C121. The structure was determined by the molecular replacement using the Phaser module in the Phenix program^23^. The searching of the initial phase was performed by using the motavizumab structure (PDB 3IXT) and the partial AF2 model of RSVF-multi-1 as a search model. Manual model building was performed using Coot^24^, and automated refinement using Phenix Refine. The final refinement statistics are summarized in Supplementary <u>Table 2</u>.

#### X-ray crystallization and structural determination of RSVF-multi-4

##### Crystallization

The RSVF-multi-4 immunogen was purified by size exclusion chromatography using a Superdex200 26 600 (GE Healthcare) equilibrated in 10 mM HEPES pH 7.5 and 150 mM NaCl and concentrated to ∼20 mg/ml (Amicon Ultra-15, MWCO 3,000). Crystals were grown at 18 C using the sitting-drop vapor-diffusion method in drops containing 1 μl of protein complex and 1 μl of reservoir solution containing 0.2 M calcium acetate, Tris pH 7.5, and 25 % (v/v) PEG 2000 MME.

##### Data collection and structural determination

Diffraction data were recorded with id30a1 at the European Synchrotron Radiation Facility, Switzerland. The diffraction data were integrated and processed to 2.382 Å by AutoProc with a high-resolution cut at I/σ=1 applied. The crystals belonged to space group P 21 21 21. The structure was determined by the molecular replacement using the Phaser module in the Phenix^23^ program. The searching of the initial phase was performed by using AF2 model of RSVF-multi-4 as a search model. Manual model building was performed using Coot^24^, and automated refinement using Phenix Refine. Unresolved loops were not built. The final refinement statistics are summarized in Supplementary <u>Table 2</u>.

#### Structure and sequence space analysis

##### Yeast display structural and sequence space analysis

For analyzing the structure space of proteins from the RSVFV yeast library, we calculated an all-vs-all TM-score^25^. This allowed us to assess the structural similarity of the designs belonging to 444 backbone families and to visualize this structural space with MDS. Additionally, we calculated the success rate of each backbone family by calculating the fraction of sequences within each family that were identified in the binding population. The five most successful families were defined as the backbone families consisting of the largest fraction of successful sequences. All sequences within each of the five most successful families were scored for sequence identity to each other using MMseqs2^15^.

##### CATH structural space analysis

In order to assess the novelty of the 8 characterized proteins and their relationship to known topologies, we compared their structural similarity to the CATH database^26^. The CATH dataset partitions proteins into a structural hierarchy; class, architecture, topology and homologous superfamily. To visualize the topological similarity space we used MDS to embed into two dimensions the matrix of all pairwise TMscores^25^ of the union of our 8 immunogens and CATH representatives from each of the 1472 known topologies. To further assess the novelty, we created two reference distributions of TM-scores against CATH; one for proteins being sampled with a novel topology and another for proteins sampled with a topology that commonly occurs in the dataset. The novel distribution was achieved by removing a topology from CATH and calculating the maximum TM-score between any protein in the excluded topology to all proteins from other topology families. The non-unique distribution was achieved by repeating this process without removing the topology. The third dataset was created by comparing our 8 designs with CATH. We compared the maximum TM-scores from our characterized proteins to these reference distributions to identify structurally similar topology families as a proxy for novelty, as previously described^14^.

