## supplementary_materials for "Accurate single domain scaffolding of three non-overlapping protein epitopes using deep learning"

### Supplementary Figures

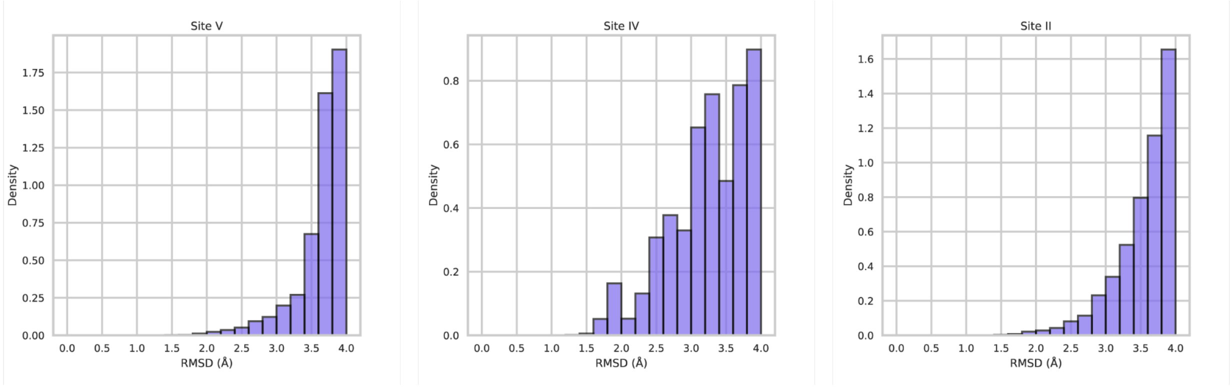


#### Supplementary Figure 1: Available natural scaffolds to host RSVF epitopes

**A-C)** Number of template structures in the PDB that can accommodate each motif targeted for scaffolding. A MASTER search was performed over a non-redundant PDB list containing a total of 8133 structures. The count of the structures recovered is plotted on the y-axis.

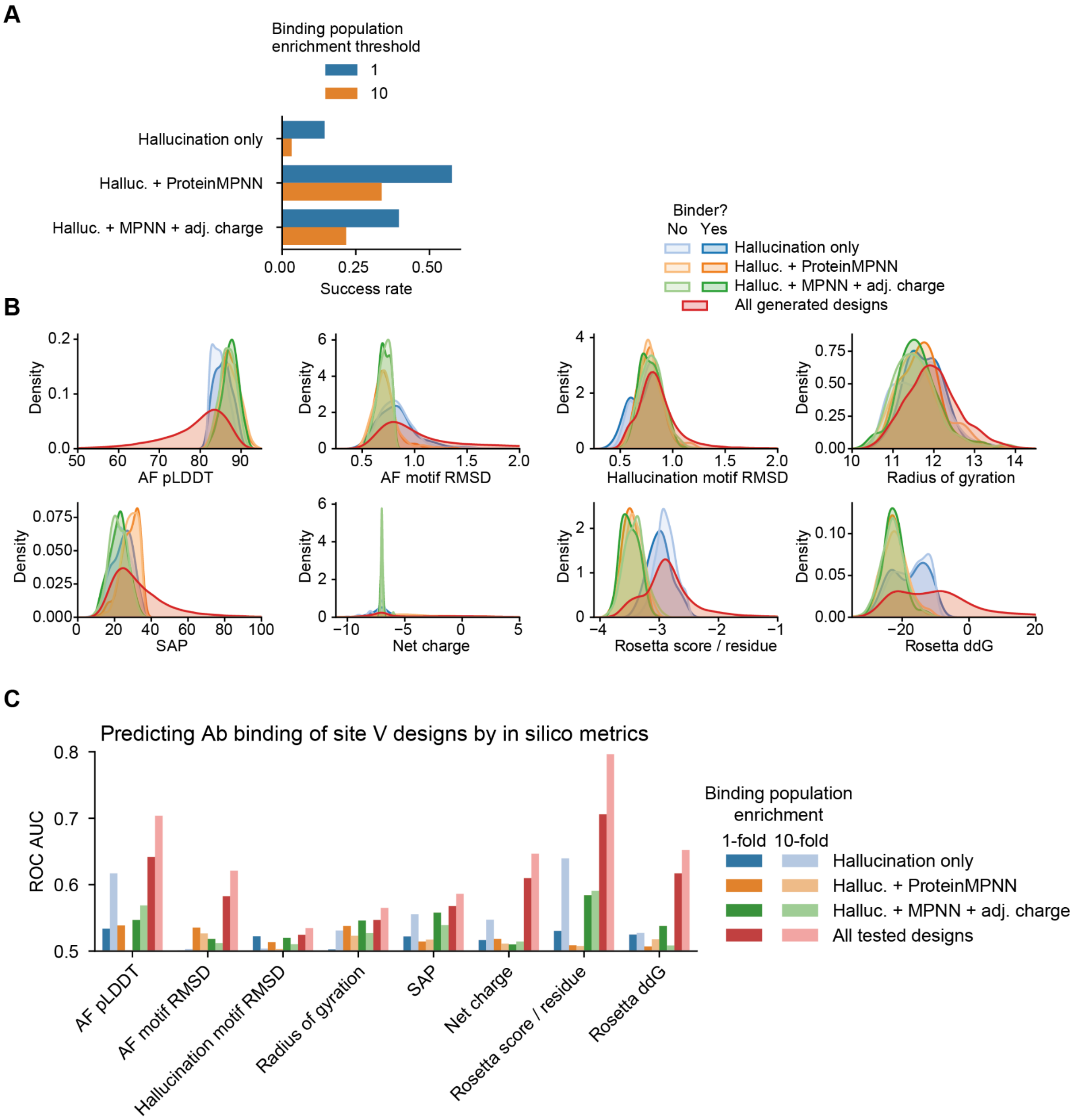


Supplementary Figure 2. Pooled analysis of RSVF site V designed binders

**A)** Success rates of RSVFV designs from the 3 design pipelines. **B)** Distributions of all *in silico* metrics used to filter designs prior to ordering for synthesis experimental testing. **C)** Receiver-operating-characteristic area under the curve for predicting binding success using each of the filtering metrics.

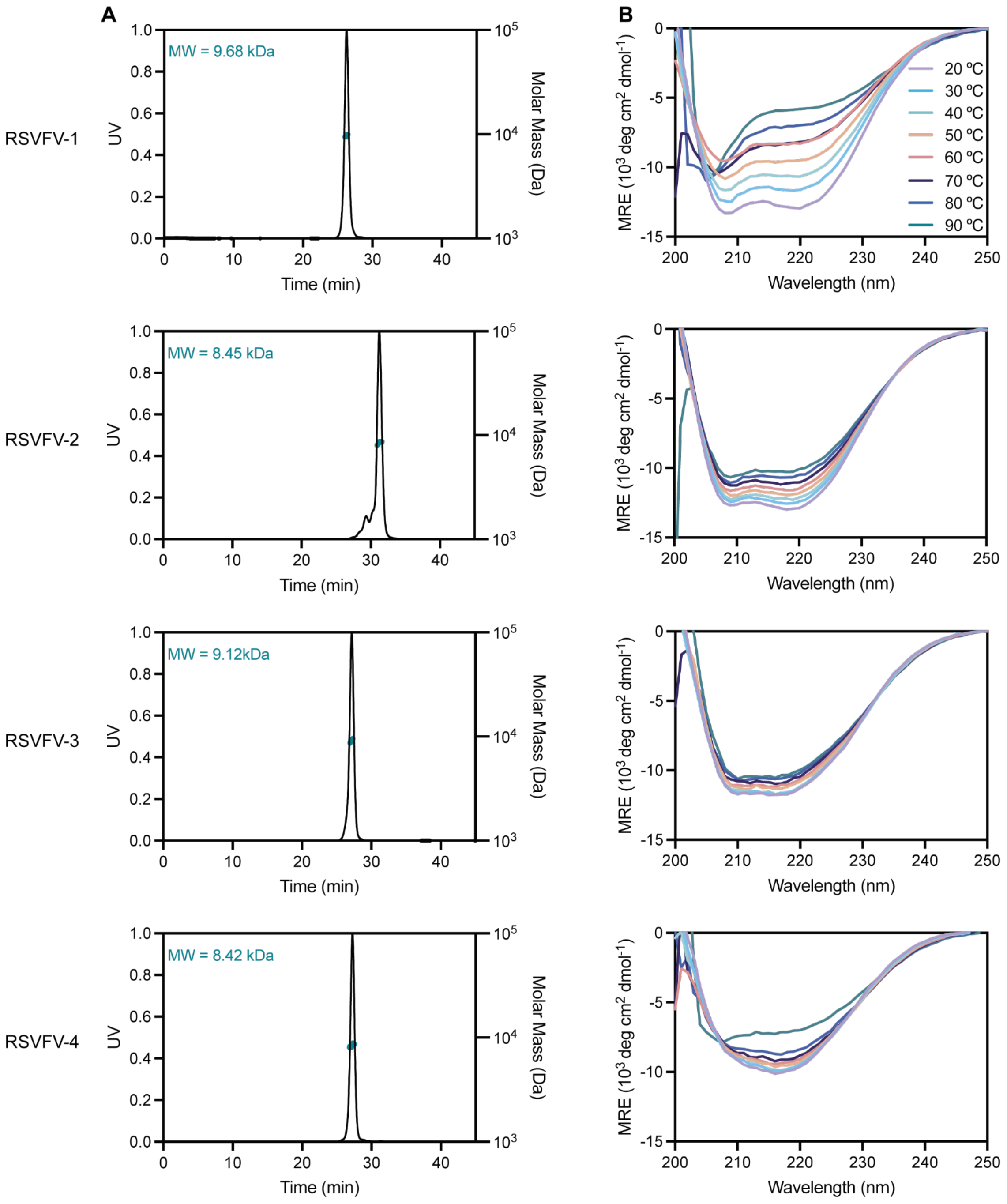


Supplementary Figure 3: RSVFV biochemical characterization

**A**) SEC-MALS measurement of oligomerization for each top candidate scaffold **C**) CD spectra at various incubation temperatures shown for each scaffold.

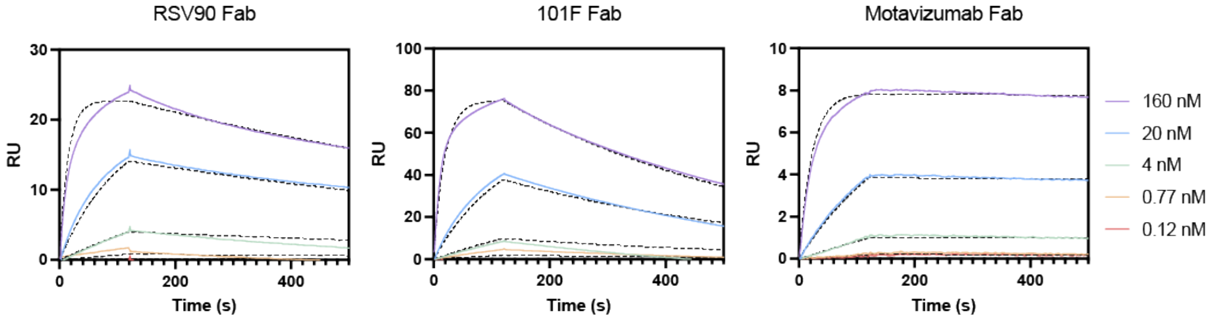


Supplementary Figure 4: RSVF trimer affinities

SPR affinity measurement of immobilized RSVF trimer against RSV90 Fab, 101F Fab, or Motazivumab Fab. Kinetics were fit using a 1:1 Langmuir model. Affinities to RSVF trimer: RSV90 Fab 0.9 nM, 101F Fab 2 nM, motavizumab Fab 15 pM

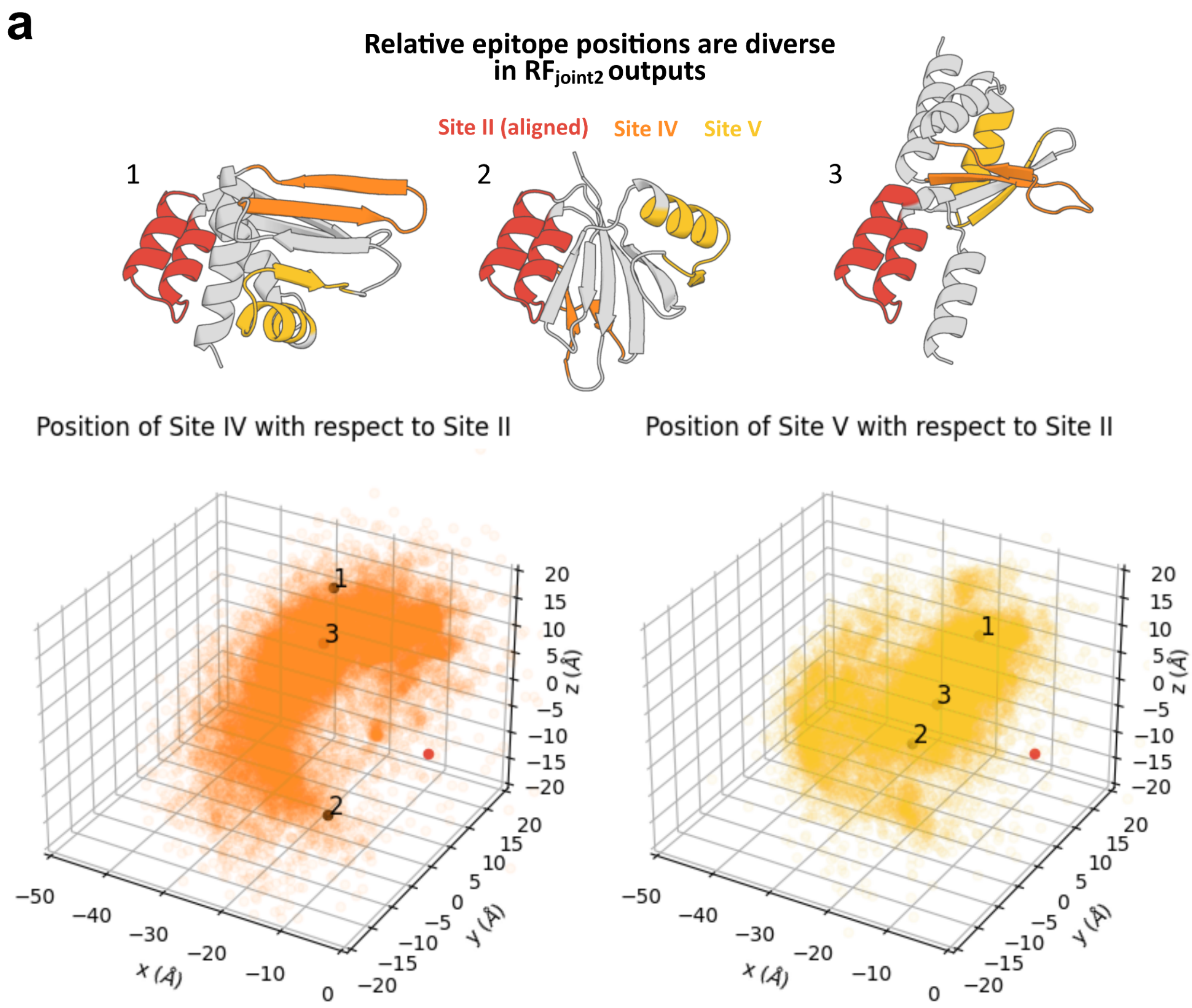


Supplementary Figure 5: RFjoint2 generates diverse inter-epitope positioning

**A)** As the relative position between epitopes is not fixed in RFjoint2, the multi-epitope designs have diverse inter-epitope positioning. Top row: three example *in silico* successful designs aligned to RSV-F Site II (red). Note the variation in the positioning of the Site IV and Site V epitopes (orange, yellow). Bottom row: quantification of the diversity of epitope positioning. Designs were aligned on RSV-F Site II, and the vector from the Site II center of mass (COM) to the COM of the other two epitopes was calculated. Red point indicates the COM of the aligned RSV-F Site II epitope in each design.

A

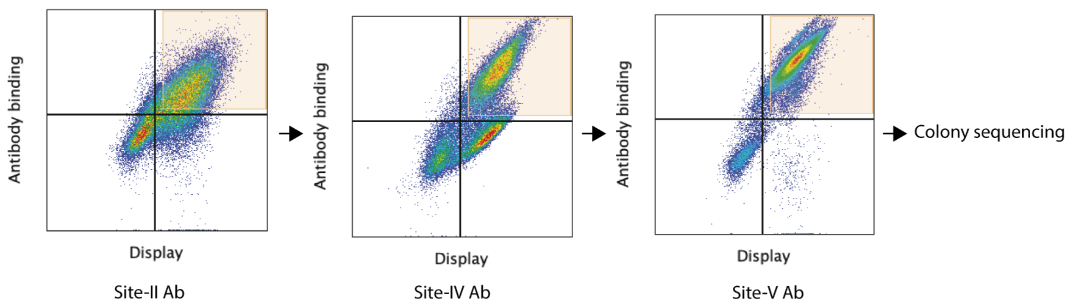


Supplementary Figure 6: Sequential sorting for the identification of multi-epitope scaffolds binding all target antibodies.

**A)** RSVF multi-epitope scaffold library screening by yeast display. Density plot of the library of scaffolds binding motavizumab antibody (site-II-specific) following proteolytic treatment. The binding population was sorted for subsequent screening against 101F antibody (site-IV-specific). The binding population was sorted for screening against RSV90 antibody (site-V-specific). The sorted population (orange square) is shown for each plot. Collected colonies binding all three antibodies were sequenced.

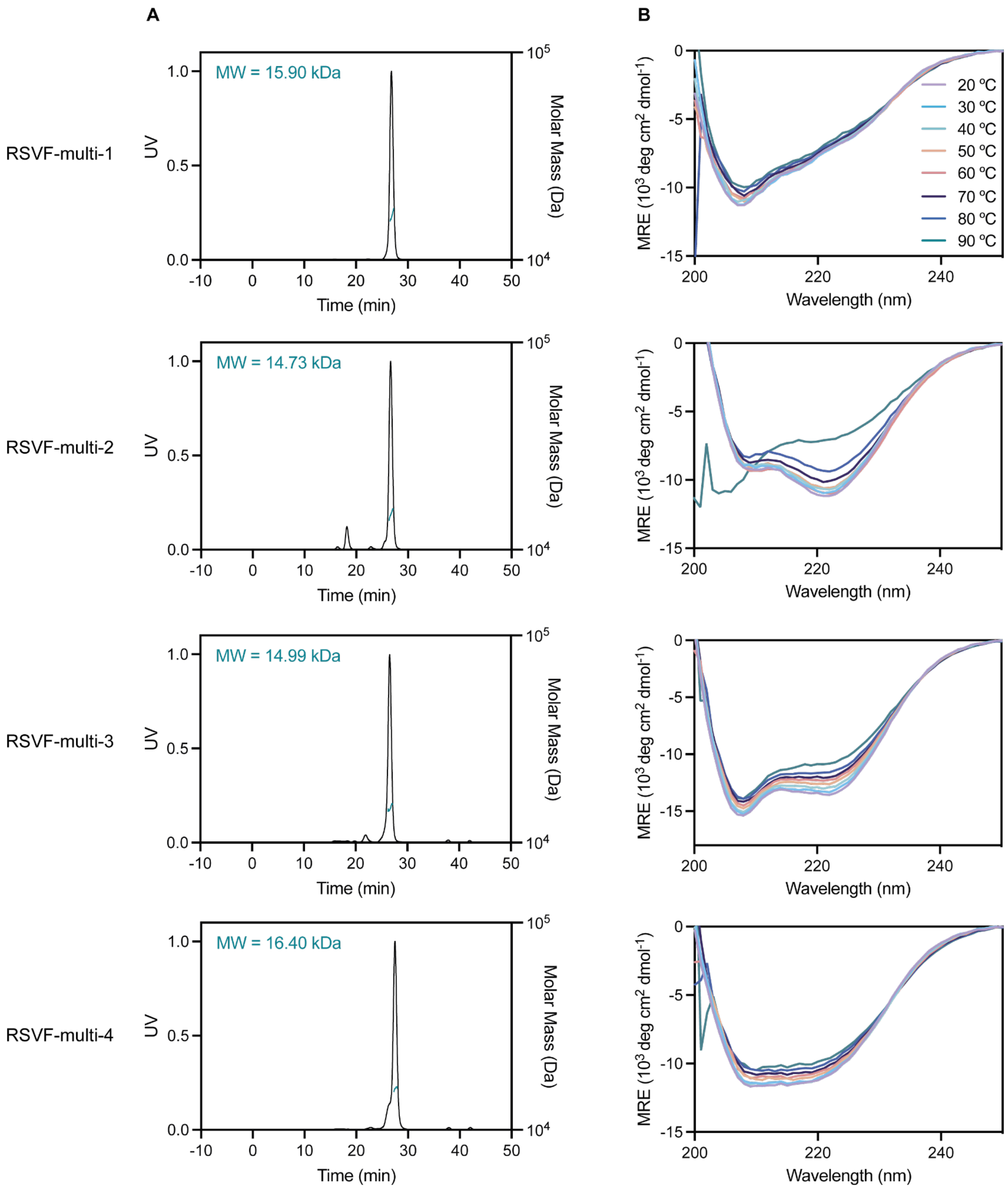


Supplementary Figure 7: RSVF-multi biochemical characterization

**A**) SEC-MALS measurement of oligomerization for each top candidate scaffold **C**) CD spectra at various incubation temperatures shown for each scaffold.

Supplementary Table 1: Affinity RSVFV and RSVF-multi designs site-specific antibodies

| | **RSV90 Fab** | **101F IgG** | **Motavizumab IgG** | **RSV90 IgG** |
|---|---|---|---|---|
| **RSVFV-1** | 54 nM | | | |
| **RSVFV-2** | 108 nM | | | |
| **RSVFV-3** | 94 nM | | | |
| **RSVFV-4** | 241 nM | | | |
| **RSVF-multi-1** | | 522 nM | – | > 10uM |
| **RSVF-multi-2** | | 377 nM | 47 nM | > 10uM |
| **RSVF-multi-3** | | 343 nM | 18 nM | > 10uM |
| **RSVF-multi-4** | | 890 nM | 14 nM | > 10uM |

Supplementary Table 2: Structural accuracy of grafted RSVF epitopes on single and multi-motif scaffolds.

The RMSD of $C_\alpha$, C, N, CO, and $C_\beta$ atoms of grafted RSVF epitopes on scaffolds compared to crystal structures of RSVF native epitopes (PDB:5TPN).

| Crystal Structure | RMSD (Å) Site-II | RMSD (Å) Site-V | RMSD (Å) Site-IV |
|---|---|---|---|
| RSVFV-1 | | 0.843 | |
| RSVF-multi-1 | 0.536 | 1.776 | 1.713 |
| RSVF-multi-4 | 0.393 | 0.790 | 1.355 |

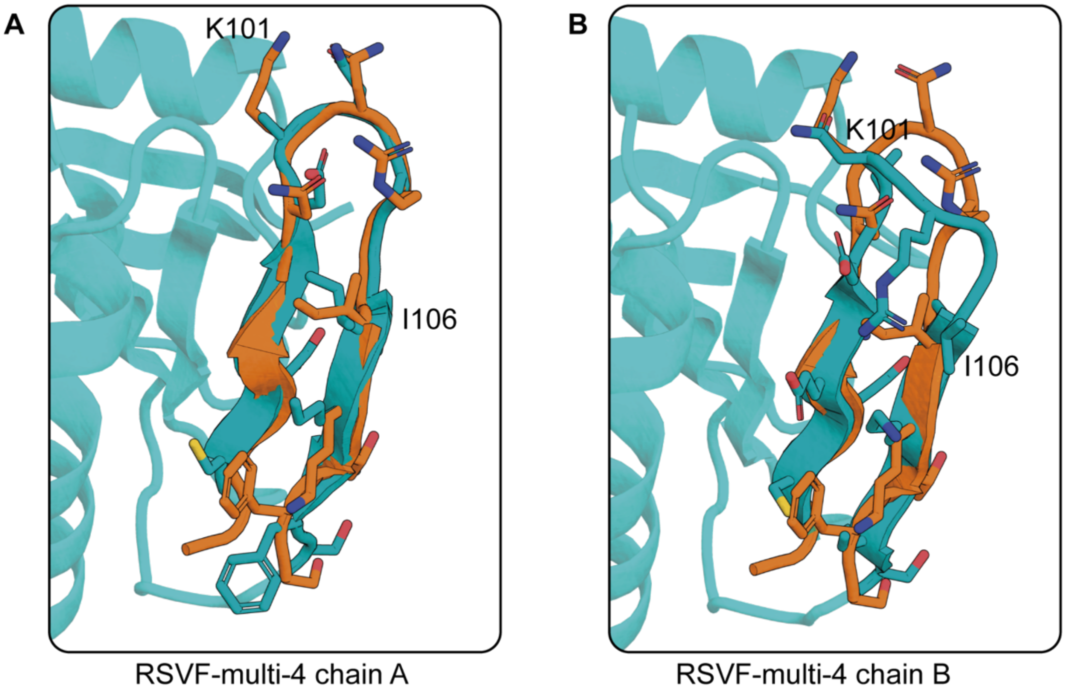


Supplementary Figure 9: Site-IV variability in RSVF-multi-4 crystal structure

Native site-IV (orange) (PDB: 3O45) overlaid on the crystal structure of RSVF-multi-4 (teal) A) Chain A residues 101-106 closely agree with the native epitope backbone. B) Chain B residues 101-106 deviate from the native epitope backbone.

Supplementary Table 3: Crystallography collection and refinement statistics:

X-ray data collection and refinement statistics for RSVF-V and multi-epitope scaffold alone or in complex with their target antibodies.

| | **RSVF-multi-4** | **Motavizumab fab/RSVF-multi-1** | **RSV90 Fab/RSVF-V H6** |
|---|---|---|---|
| Wavelength | 0.96 | 0.87 | 1.00 |
| Resolution range | 77.67 - 2.3 (2.382 - 2.3) | 78.23 - 2.91 (3.014 - 2.91) | 73.83 - 2.43 (2.517 - 2.43) |
| Space group | P 21 21 21 | C 1 2 1 | P 21 21 2 |
| Unit cell | 46.843 106.432 113.589 90 90 90 | 184.072 66.859 109.41 90 103.865 90 | 113.39 137.65 87.47 90 90 90 |
| Total reflections | 135054 (13569) | 86269 (8958) | 332044 (34355) |
| Unique reflections | 25945 (2558) | 28467 (2829) | 52204 (5153) |

| | | | |
|---|---|---|---|
| Multiplicity | 5.2 (5.3) | 3.0 (3.2) | 6.4 (6.7) |
| Completeness (%) | 99.26 (99.96) | 99.67 (99.82) | 99.79 (99.96) |
| Mean I/sigma(I) | 9.17 (1.55) | 6.14 (1.52) | 9.58 (1.44) |
| Wilson B-factor | 54.13 | 45.54 | 52.47 |
| R-merge | 0.08944 (0.9056) | 0.1444 (0.7102) | 0.1489 (2.189) |
| R-meas | 0.09991 (1.005) | 0.1753 (0.8547) | 0.1623 (2.377) |
| R-pim | 0.04371 (0.4287) | 0.09815 (0.4709) | 0.06398 (0.9186) |
| CC1/2 | 0.996 (0.77) | 0.989 (0.738) | 0.997 (0.58) |
| CC* | 0.999 (0.933) | 0.997 (0.921) | 0.999 (0.857) |
| Reflections used in refinement | 25819 (2558) | 28586 (2828) | 52183 (5152) |
| Reflections used for R-free | 1357 (132) | 1407 (142) | 2632 (243) |
| R-work | 0.2658 (0.3414) | 0.2228 (0.3267) | 0.2462 (0.4039) |
| R-free | 0.2962 (0.3935) | 0.2600 (0.3622) | 0.2866 (0.4492) |
| CC(work) | 0.940 (0.779) | 0.941 (0.828) | 0.946 (0.777) |
| CC(free) | 0.923 (0.710) | 0.910 (0.757) | 0.946 (0.725) |
| Number of non-hydrogen atoms | 3968 | 8365 | 7750 |
| macromolecules | 3908 | 8316 | 7704 |
| ligands | 0 | 4 | 0 |
| solvent | 60 | 45 | 46 |
| Protein residues | 493 | 1078 | 1016 |
| RMS(bonds) | 0.005 | 0.007 | 0.002 |
| RMS(angles) | 0.89 | 1.07 | 0.51 |
| Ramachandran favored (%) | 97.09 | 96.42 | 95.91 |
| Ramachandran allowed (%) | 2.91 | 3.58 | 3.89 |
| Ramachandran outliers (%) | 0 | 0 | 0.20 |
| Rotamer outliers (%) | 0.23 | 1.17 | 0.82 |

| | | | |
|---|---|---|---|
| Clashscore | 4.15 | 9.1 | 5.54 |
| Average B-factor | 66.14 | 50.71 | 74.65 |
| macromolecules | 66.28 | 50.71 | 74.75 |

Supplementary Table 4: Sequences of experimentally characterized designs

| Name | Sequence | Expression vector |
|---|---|---|
| RSVFV-1 | METEEEIIEKVKSALLSTNKAVISVELKGRTIPLYVEITKEGKL HLTAEGATEEEKEIIKEAQKAFQEEIEHEAERKEK | pet11b |
| RSVFV-2 | SAELDVKAEAIVNKIESALLSTNKAVVSWEGKTLTVTLENNT LIIEVEEVDEEMKELLEKAAKLWEEKKGKKAEVILP | pet11b |
| RSVFV-3 | MVTKEEIINKIKSALLSTNKAVVSIKNPKTNEYVPFLVTNNGG EIVVEDTNGNKFVSKNSLEDVANWILEYLK | pet11b |
| RSVFV-4 | MTPEEAKELYEKAKSALLSTNKAVISAEINGKTLTAEVSLTS DNKIEVTITEGDKTTTITFDTNDENKYTTEETSA | pet11b |
| RSVF-multi-1 | MKLVIARVKSPKVKRLSEEDIEKIKSALKSTNKAVVTIKDENG EEIEVEVRLLTLEEALKYINDLPISNDAKKLMSNNIHKALEPG RTVVFGPEGCEERDKNRGIIKTFSTDVKLDETYFFFRVE | pet11b |
| RSVF-multi-2 | MKLVIARVKSPKVKRLSEEDIEKIKSALKSTNKAVVTIKDENG EEIEVEVRLLTLEEALKYINDLPISNDAKKLMSNNIHKALEPG RTVVFGPEGCEERDKNRGIIKTFSTDVKLDETYFFFRVE | pet11b |
| RSVF-multi-3 | DDLVDIFLRAFAKAAKVTRFDKNRGIIKTFSEEETEKLFKSLT EEEVEKIESALKSTNKAVVVLGDGDIEIEIDLDKLYALINDLDI SNDQKKEMSNNFFEYLRKIAKK | pet11b |

| | | |
|---|---|---|
| RSVF-multi-4 | MKLVIARVKSPKVKRLSEEDIEKIKSALKSTNKAVVTIKDENG EEIEVEVRLLTLEEALKYINDLPISNDAKKLMSNNIHKALEPG RTVVFGPEGCEERDKNRGIIKTFSTDVKLDETYFFFRVE | pet11b |
